# Spatiotemporal conservation of bacteria in the *Trichodesmium* microbiome

**DOI:** 10.64898/2026.08.28.744692

**Authors:** Anjali M. Bhatnagar, Nina Yang, Catie S. Cleveland, Shelby J. Barnes, Eric A. Webb

## Abstract

*Trichodesmium* is a major nitrogen fixer in the nutrient-limited tropical and subtropical ocean, contributing up to 50% of new nitrogen and driving primary productivity. *Trichodesmium* filaments often aggregate into colonies that host a diverse consortium of microbes that can influence their activity and physiology; however, it is unclear what factors shape community structure. To address this, we used metagenomics of *Trichodesmium* colonies collected across the subtropical and tropical Atlantic Ocean to examine how *Trichodesmium* diversity and abundance shape epibiont community composition. Heterotrophic metagenome-assembled-genomes from *Trichodesmium* colonies and genes specific to the major *Trichodesmium* subclades were read mapped against *Trichodesmium* metagenomes from the North Atlantic, the Western Tropical South Pacific, and the Red Sea. Using this global survey of *Trichodesmium* subclade distribution patterns and their enriched heterotrophs, we found unique core microbiomes that are specific to *Trichodesmium* subclades across ocean basins. Genome characteristics of the enriched heterotrophs reveal a genomically small, slow-growing community consistently associates with colonies. Our results suggest that *Trichodesmium* genotype may influence epibiont community structure, indicating a role for deterministic processes in shaping the microbiome.

## 2. Introduction

Nitrogen (N) constrains phytoplankton growth and biogeochemical cycling across much of the oligotrophic ocean (1–5). In these regions, which span nearly 40% of the world’s oceans (6), diazotrophs convert inert N_2_ into biologically available N (4,7,8), driving primary productivity and the biological carbon (C) pump (8). Among them, the filamentous cyanobacterium *Trichodesmium* is one of the most important diazotrophs, contributing up to half of global marine N_2_ fixation (1–4). Consequently, a large body of research has focused on understanding the environmental controls on *Trichodesmium* physiology and biogeography, particularly nutrient availability, including iron (Fe) and phosphorus (P) (4,8–18). However, growing evidence suggests that biotic interactions, especially with the microbiome, also influence *Trichodesmium* growth and N_2_ fixation.

In the open ocean, individual *Trichodesmium* filaments aggregate into visible (> millimeter-sized) colonies that provide a stable habitat for diverse epibiotic microorganisms (19). These microbiome members are hypothesized to increase the bioavailability of growth-limiting nutrients, such as P and Fe, to benefit *Trichodesmium* (20,21). Consistent with this, *Trichodesmium* grows approximately twice as fast in the presence of heterotrophs than axenically (22,23), suggesting that the microbiome contributes to *Trichodesmium’*s success in the oligotrophic ocean (20,21,24).

Metagenomic investigations of *Trichodesmium* colonies reveal epibiotic microorganisms that recur across seasons (25) and ocean basins (26). This “core” microbiome is often supplemented by a variable community of epibionts that changes with geography or environmental conditions (25,27). However, the processes governing microbiome assembly remain poorly understood. In phytoplankton, microbiome composition is shaped by both deterministic (selection imposed by abiotic or biotic factors) and stochastic (random) processes, though their relative influence varies among hosts (28–30). Deterministic assembly may reflect environmental filtering (30) or host genetic background (31), which together determine which microbial taxa persist within the community.

The extent to which deterministic selection by the environment or *Trichodesmium* genetic diversity, versus stochastic processes, shapes microbiome composition and stability remain poorly understood. Natural *Trichodesmium* communities are mixes of the two dominant clades, *Trichodesmium erythraeum* (Tery) and *Trichodesmium thiebautii* (Thieb) (9,32–37), however, Thieb is the numerically dominant clade present (34). Recent phylogenomics efforts with new enrichment cultures further resolve Tery and Thieb into the TeryA, TeryB, ThiebA, ThiebB, ThiebC, and ThiebD subclades (38,39). Among these, ThiebD is the most abundant open-ocean subclade (38,39). Variation in the distribution of these subclades may be linked to differences in microbiome compositions that differentially influence *Trichodesmium* growth and N_2_ fixation through nutrient exchange under fluctuating environmental conditions (20,24,26,40,41). Given *Trichodesmium*’s major contribution to global N_2_ fixation, understanding the composition, function, and assembly of its microbiome is essential for predicting its role in marine biogeochemical cycling in a changing ocean.

In this study, we collected *Trichodesmium* colonies from the Bermuda Atlantic Time-series Study (BATS) and North Atlantic Ocean (NATL) over two summers and compared their metagenomes with published datasets from the NATL (26,37), western tropical South Pacific Ocean (WTSP) (42), and Red Sea (RS) (25,43) to investigate the processes governing microbiome assembly. Comparative genomics of metagenome-assembled genomes (MAGs) revealed a microbiome enriched in novel, genomically small, slow-growing heterotrophs. Across ocean basins, *Trichodesmium* populations remained regionally stable, and in the NATL, the dominant ThiebD subclade consistently co-occurred with a distinct set of MAGs, supporting the existence of a subclade-specific microbiome. Collectively, these findings indicate that *Trichodesmium* and its microbiome form a stable, non-random association, shaped by host population structure.

## 3. Methods

### 3.1 Sample Collection

*Trichodesmium* colonies were collected during six R/V Atlantic Explorer BATS cruises in the summers of 2022 and 2023 (**Table S1, Figure S1**), using methods previously described (39). Colonies were rinsed 3x with locally collected, sterile filtered (0.2 µm) seawater to capture epibionts that are tightly attached to or entrained in colonies. For the 2018 Station 17 (PRJNA828267), the entire contents of the net tow (>100 µm) were filtered onto 25mm 5-µm PC filter (37), representing the particle fraction in the NATL.

### 3.2 DNA and Sequencing

DNA was extracted using a modified CTAB/phenol/chloroform protocol (44). Colonies were gently washed from filters with 600 µL of TE buffer, lysed with 30 µL of 25% SDS and proteinase K (20 mg/mL, final 1.2%; Sigma-Aldrich), and incubated for 24 hours at 65°C. Filters were stored on ice until proteinase K was added. Filters were then removed, and 100 µL 5M NaCl and 100 µL CTAB (2% w/v) were added prior to extraction. All microcentrifuge steps were performed at room temperature. DNA quantity and quality were assessed using a NanoDrop Spectrophotometer (Thermo Scientific, MA, USA).

Samples with A260/A280 ratios between 1.8–2 were submitted to Novogene (Sacramento, CA, USA) for microbial whole-genome metagenomic sequencing on the NovaSeq 6000 platform (Illumina) using 150 bp paired-end reads, targeting 25 Gbps per sample.

### 3.3 Ocean-basin Meta-analyses for *Trichodesmium*

Total *Trichodesmium* abundance in each sample was estimated by read-recruitment to representative genomes from the Tery (IMS101) and Thieb (H94) clades across 46 metagenomes in the NATL (this study and (26,37)), WTSP (42), and RS (25,43)(**Table S1**). Samples with low *Trichodesmium* content (< 15%) were removed from downstream analyses, except for Station 17, which represents a bulk phytoplankton sample > 100 µm (**Table S1**).

Because *Trichodesmium* populations are difficult to resolve using 16S rRNA gene and ITS sequence alone (33), fine-scale diversity was assessed by read-recruitment to conserved, subclade-specific genes from TeryA, TeryB, and ThiebA-D (**Table S2**) in all 46 metagenomes (**Figure S2**) using the Anvi’o v8 read-recruitment workflow (45). Genes identified in previous studies (38,39) were concatenated into one FASTA file, and BAM files were filtered with CoverM v0.6.1 (46) to retains reads matching at 99.5% identity along 90% of the read. Relative subclade abundance was calculated from the average depth of coverage (Q2Q3), which normalizes for differences in sequencing depth and gene length.

### 3.4 Metagenome-Assembled Genome (MAG) Assembly

Metagenome-assembled genomes (MAGs) were assembled as previously described (39) with modifications (**Supplementary Methods**). Briefly, reads were assembled into contigs using MEGAHIT (metasensitive preset) (47), and binned MaxBin2 v2.2.4 (48), MetaBAT2 v1.7 (49), and CONCOCT v1.1 (50). To accurately compare between datasets, MAGs from the WTSP (42) and RS were reassembled using the methods in this paper; reassembled MAGs were similar to those described in (25). *Trichodesmium* MAGs were refined at ≥ 70% completion and < 5% contamination. Heterotroph MAGs were refined at > 50% completion and < 5% contamination (**Table S4**).

### 3.5 Taxonomic Placement

New and published MAGs were phylogenomically characterized as previously described (37)(**Table S3-Table S4**). *Trichodesmium* MAGs from BATS were placed alongside 16 published *Trichodesmium* MAGs and isolate genomes using 251 core cyanobacterial genes (51), with *Okeania* (GCF010692555.1; GCF010672385.1) as the outgroup. Heterotroph MAGs (**Table S4**) were placed using 74 core bacterial genes, with green sulfur bacteria (GCF000281175.1; GCF000021945.1;GCF000017805.1) as the root (52). An Alphaproteobacteria-specific phylogeny was constructed using genomes and isolates in (53–60) along with GCA_000419545.1, SAR11 MED-G05 and SAR11 MED-G06. *E. coli* (GCA 001544635.1) served as the outgroup. Trees were visualized and annotated in iTol using the annotation function (61).

### 3.6 Ocean-basin Meta-analyses for Assembled Heterotrophic MAGs

The Anvi’o v8 (45) read-recruitment workflow was used to determine bacterial epibiont distributions across the 46 metagenomes, with CoverM filtering reads at ≥ 95% identity along 99% of the read. Samples with fewer than 10 million reads were removed to ensure epibiont detection (**Table S1**). To avoid nonspecific read recruitment, a representative, MAG (≥ 70% complete, < 5% contamination) was picked for read-recruitment based on GTBD-tk classification, phylogenetic placement, and ANI following dereplication at 98% ANI (**Table S4**). *Alteromonas macleodii* was represented by GCF_002849875.1. MAGs were considered present if ≥ 40% of the genome was detected. Statistical analyses were all done in Rstudio v4.0.2 using the vegan package (62) and heatmaps were made using ComplexHeatmap v2.13.1 (63). For heatmap visualization, *Trichodesmium* populations were hierarchically clustered.

### 3.7 Normalized Stochastic Ratio

The normalized stochastic ratio measures the relative proportion of deterministic and stochastic effects in a community across large spatial scales (64). A normalized stochastic ratio of 0% indicates deterministic forces shape community assembly and a ratio of 100% means that stochastic effects play a stronger role. The stochastic ratio was calculated using the NST package v3.1.10 in R (65). Due to the sparse nature of the presence/absence dataset, percent genome detection values (**Supplementary Methods**) were Hellinger-transformed and Euclidean distance was used as the dissimilarity metric. Values were not included for the WTSP due to insufficient samples.

### 3.8 Indicator Species Analysis and Heterotroph Correlations

Indicator species analysis (ISA) was used to identify heterotrophs associated with specific *Trichodesmium* populations and sampling locations (66,67). Analyses were conducted on heterotroph presence/absence data using the ‘indicspecies’ package v1.8.0 (68). Because *Trichodesmium* colonies often contain multiple, co-occuring subclades, samples were grouped according to their relative subclade abundance. Indicator associations with a single subclade (e.g., ThiebD) reflect enrichment in colonies dominated by that subclade, whereas associations with subclade combinations (e.g., TeryB-ThiebD or ThiebD-ThiebC) reflect enrichment in colonies containing co-dominant populations.

Associations between heterotroph correlations and *Trichodesmium* subclades were further evaluated using linear correlations between heterotroph percent genome detection and the average coverage of subclade-specific genes. Due to the mixed nature of colonies, correlations were restricted to samples dominated by a single subclade. Statistical significance is reported as false discovery rate (FDR)-adjusted p-values and visualized as -log10(FDR-adjusted p-value).

### 3.9 Predicted Growth Rate and Genomic Potential of the Heterotrophic MAGs

Genome-wide codon usage statistics generated with gRodon v2.4.0 (69) were used to estimate the minimum hourly doubling time of 105 ‘enriched’ heterotrophic genomes. Genomic potential was determined through gene prediction and annotation using Prodigal v2.6.3 (70), PROKKA v1.13.3 (71), KEGG Pathway Annotation v1.3.0 (72) and HMMER v3.1b2 (73,74). KEGG-Decoder v0.7 (75) and Anvi’o v8 (45) were used to visualize the percent completion of gene-cluster completeness.

Genes involved in oxidative stress response (*soda*, *sodB, and katG*), phosphorus acquisition (*phoA*, *phoD*, *phoX*), and iron storage (*bfrA* and *bfrB*) were identified by BLAST searches against the Uniprot database (76). Hits were retained if they met high-confidence thresholds (evalue ≤ 1e-5, bitscore ≤ 50). Gene coverage was calculated as the proportion of the reference sequence aligned to each MAG and normalized to a 0–1 scale, where a score of 1 indicates complete coverage and 0 indicates no coverage.

## 4.0 Results

### 4.1 *Trichodesmium*-associated microbial diversity

To investigate relationships between *Trichodesmium* and its microbiome, we sequenced hand-picked colonies from BATS and the NATL in June–September of 2022 and 2023 (**Figure S1; Table S1**). To place these samples in a broader context, we analyzed them alongside published datasets in the NATL (26,37), RS (25,43), and WTSP (42), generating a global dataset of 46 metagenomes spanning multiple ocean basins and years (**Figure S2; Table S1**). From these samples, we assembled and hand-refined a total of 213 heterotrophic MAGs (completeness ≥ 50%, contamination < 5%) from all metagenomes (**Figure 1**; **Table S4**). Putative epibiont 16S rRNA sequences were classified as eukaryotes, archaea, bacteria, and other cyanobacteria (**Table S5**); however subsequent analyses focused on heterotrophic bacteria, which constituted approximately 90% of the recovered MAGs.

**Figure 1.**
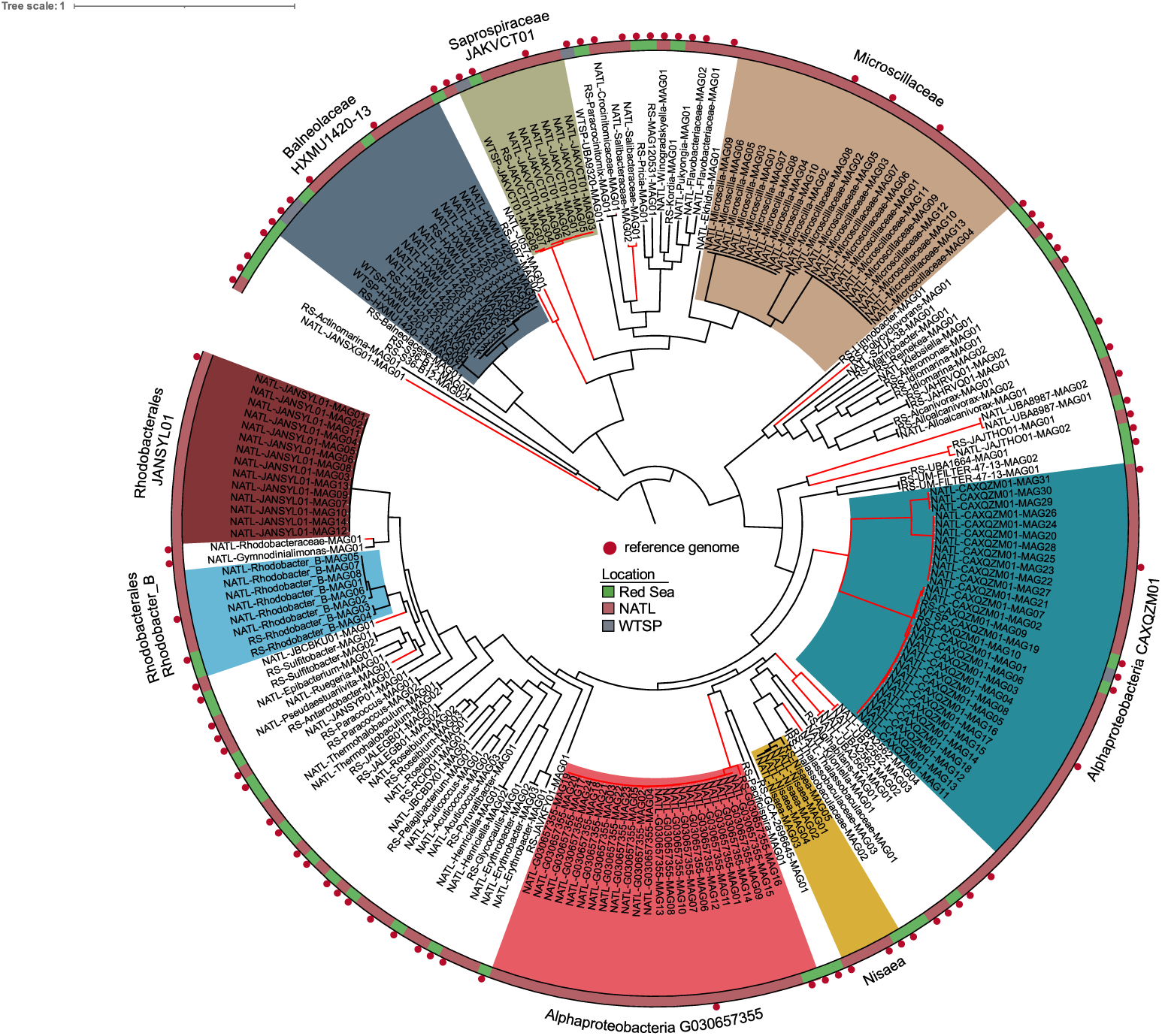
Maximum likelihood tree of 213 hand-refined MAGs assembled from handpicked *Trichodesmium* colonies from the North Atlantic (NATL), the western tropical South Pacific (WTSP), and the Red Sea (RS) with 51 conserved bacterial genes. Locations are indicated on the outside ring of the tree. Red branches show novel species as indicated by the GTDB-tk classification. Colored groups with taxonomic labels represent MAGs assembled from different samples multiple times. Red circles indicate the 83 genomes picked for read recruitment.

At a broad taxonomic level, the MAGs were dominated by Alphaproteobacteria, Bacteroidia, and Gammaproteobacteria (**Figure 1; Table S4**). Dereplication at 98% ANI yielded 83 putative epibiont species represented by a single reference MAG per taxonomic group (**Figure 1**; **Table S4**). Of these, 31% were classified as GTDB-tk red novel (77) and 75% lack species classifications or fell outside established bacterial species boundaries (**Figure 1; Table S4**).

One of the most prevalent groups, the Alphaproteobacterial family CAXQZM01, was classified as GTDB-tk RED-novel and consistently assembled in *Trichodesmium* metagenomes collected at BATS and in the NATL during 2022-2023 (**Figure 1; Table S4**). Highly similar CAXQZM01 MAGs (> 98% ANI) were also recovered from the NATL in 2018 and the RS (**Table S4**). Phylogenomic analysis placed CAXQZM01 in a highly supported cluster outside of all Alphaproteobacterial genomes available on NCBI in Fall 2022 (> 1200 MAGs; **Figure S4**). A 16S rRNA gene recovered from one CAXQZM01 MAG shared 99% identity with an Alphaproteobacterium previously isolated from *Trichodesmium* colonies in 2011 (GenBank JN790965.1; JN790966.1).

Several additional MAGs were repeatedly assembled across independent sampling times and locations (**Figure 1**), including members of Balneolaceae (*HXMU1420-13*; 16 times), Saprospiraceae (*JAKVCT01*; 8 times), Microscillaceae (23 times), G030657355 (24 times), CAXQZM01 (31 times) and the Rhodobacterales genera *JANSYL01* (14 times) and Rhodobacter_B (8 times). While Microscillaceae and *JANSYL01* were restricted to western NATL samples, the remaining taxa were recovered across multiple ocean basins. These recurrent MAGs represent enriched and persistent members of the *Trichodesmium* microbiome and are hereafter referred to as the “enriched” community.

### 4.2 Spatiotemporal Variation in *Trichodesmium* Populations and Associated Heterotrophs

Regional and global patterns of microbial community composition were assessed using a nonmetric multi-dimensional scaling (NMDS) analysis based on Jaccard matrices of heterotroph presence/absence and *Trichodesmium* populations (**see Supplemental Methods; Figure S5; Table S6)**. Within the NATL, heterotroph communities clustered into western, mid, and eastern Atlantic (R^2^ = 0.16, p value < 0.01) (**Figure 2A**). In the global dataset, heterotrophic communities formed two clusters: one containing all NATL samples and another containing samples from both the RS and WTSP (R^2^ = 0.24, p value < 0.001) (**Figure 2B**). Similar clustering patterns were observed when using percent genome detection (**Figure S4**; **see Supplemental Results and Discussion**). *Trichodesmium* in the global dataset also clustered by populations (R^2^ = 0.48, p value < 0.001) (**Figure 2C**). Statistical comparisons involving WTSP samples were limited due to low sample size.

**Figure 2.**
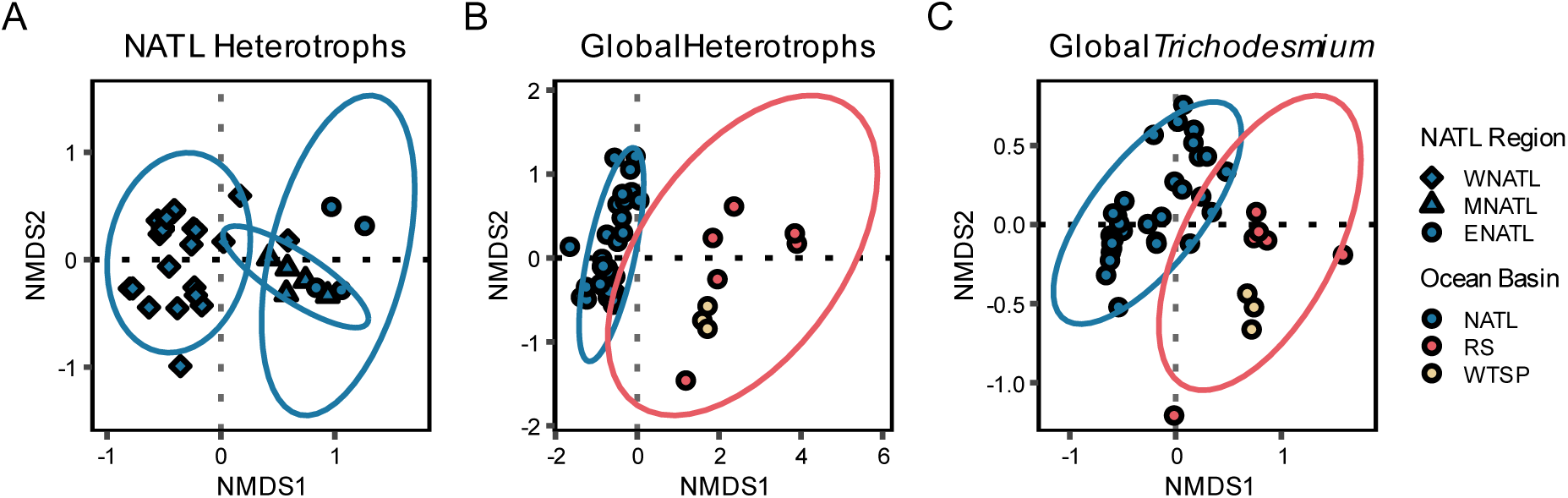
Non-metric Multidimensional Scaling (NMDS) of *Trichodesmium* and heterotroph community composition. Heterotroph presence was determined if ≥ 40% of the MAG was detected in a sample. **(A)** NMDS of the Jaccard distances of the presence/absence of heterotroph MAGs detected in the NATL (3D Stress = 0.075) **(B)** NMDS of the Jaccard distances of the presence/absence of heterotroph MAGs detected in the NATL, RS, NPSG, and WTSP (3D Stress = 0.056) **(C)** NMDS of the normalized Bray-Curtis dissimilarities between *Trichodesmium* communities in the NATL, RS, NPSG, and WTSP (3D Stress = 0.047). Colored ellipses show a 95% confidence interval for samples in the North Atlantic (blue) and the Red Sea (red). Statistical significance was not calculated for the WTSP due to lack of samples.

We applied the normalized stochasticity ratio to the Euclidean distances of the Hellinger-transformed heterotroph percent genome detection matrix (NST_euc_) (64) to assess the relative contributions of deterministic and stochastic processes to microbiome assembly (**Table S6**). The ratio in the NATL (n = 30) was 17.8% on a scale of 0–100% where 50.0% marks the boundary between deterministic and stochastic assembly processes. In the West NATL (n = 21), the ratio is 17.5%. Similar values are seen in the East NATL (n = 4) and Mid NATL (n = 5) at 17.4% and 18.7%, respectfully. The ratio in the RS was 38.0% (n = 5).

We cataloged changes in colonial *Trichodesmium* subclade composition across spatiotemporal scales. Based on read recruitment to representative Thieb and Tery genomes, *Trichodesmium* content averaged 43.01% ± 15% (**Table S1**). Subclade composition was resolved using conserved subclade-specific gene clusters (**Table S2**) (38,39), and populations were classified by the relative abundance of the primary subclade(s): single-dominant (one subclade, > 50%) co-dominant (two subclades comprising ∼50%), or mixed (three or more subclades occurred at ∼equal proportion) (**Table S7; Figure S6**). As a result, 24 samples were single dominant, 15 were codominant, and one was mixed.

*Trichodesmium* population composition varied across ocean basins (**Figure 3A**). In the NATL, colonies transitioned from ThiebD-enriched populations in the western basin to mixed ThiebD-ThiebC populations in the mid-Atlantic and ThiebC-dominated populations in the eastern NATL (**Figure 3A**). In the RS, five samples were ThiebB-enriched communities, and one was TeryB-enriched (**Figure 3A**). The WTSP had the most subclade diversity, as samples were TeryA-ThiebB codominant (**Figure 3A**). No *Trichodesmium* were detected in the bulk tow (St17), which represented the oligotrophic particle fraction in the NATL (**Figure 3A; Table S1**).

**Figure 3.**
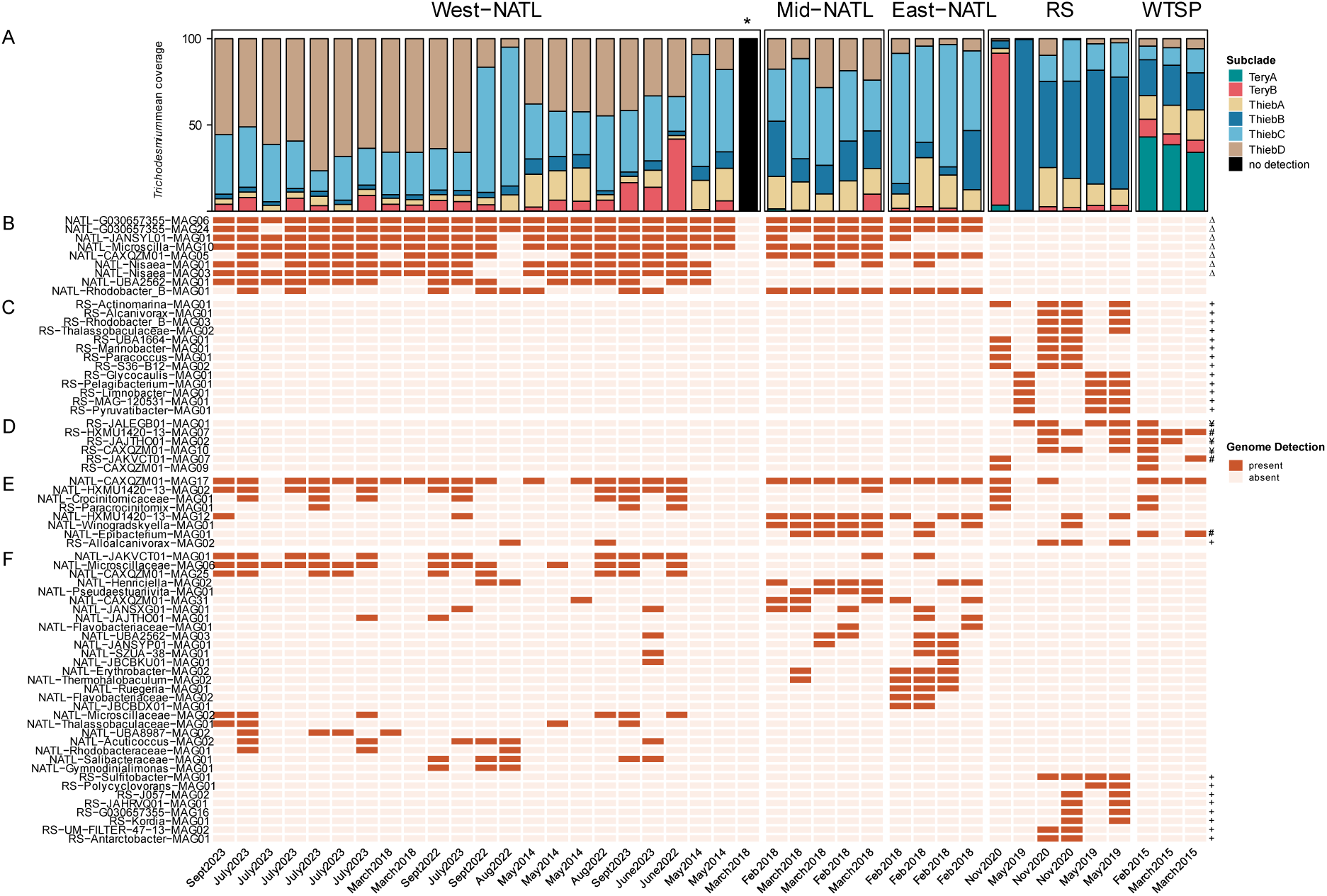
*Trichodesmium* populations and microbial genome detection (%). **(A)** Percent *Trichodesmium* subclade populations. Samples are hierarchically clustered by region. Asterisk denotes a bulk net tow (see methods). **(B-D)** MAGs were denoted as present if > 40% of their genome was detected in a sample (filled). Each row represents one MAG. **(B)** Heterotroph MAGs detected in ≥ 50% of the NATL samples. **(C)** Heterotroph MAGs detected in ≥50% of the RS samples. **(D)** Heterotroph MAGs detected in ≥ 50% of the RS and WTSP samples. **(E)** Heterotroph MAGs detected in the NATL, RS and WTSP. **(F)** Hierarchical clustering of heterotrophic MAGs. Symbols indicate MAGs identified as statistically significant in the ISA (p value < 0.05). Squares indicate MAGs associated with ThiebB and ThiebB-TeryA populations. Circles indicate MAGs associated with the following population groups: ThiebD, ThiebC, TeryB, ThiebD-ThiebC, ThiebD-TeryB, and ThiebC-ThiebB, ThiebC-ThiebA, and ThiebC-TeryB. Triangles indicate MAGs associated with the following populations: ThiebD, ThiebC, TeryB, ThiebD-ThiebC, ThiebD-TeryB, ThiebC-ThiebB, and ThiebB-TeryA samples. Sample date is labeled on the bottom. MAGs starting with ‘AE’ or ‘BVAL’ were collected from BATS for this study; MAGs starting with ‘St’ are from Webb et al., 2023; MAGs starting with ‘RS’ are from Koedoder et al., 2023 and 2022; MAGs starting with ‘WTSP’ are Frischkorn et al., 2018.

### 4.3 Identifying Conserved Heterotrophs in *Trichodesmium* Colonies Across Ocean Basins

Visually comparing *Trichodesmium* subclade composition and heterotroph MAG detection revealed distinct patterns among regions. Despite substantial variation in *Trichodesmium* subclade composition, NATL samples shared a common set of heterotroph MAGS (**Figure 3B**). In contrast, the RS and WTSP samples lacked many of these taxa but were more similar to each other, sharing five MAGs including representatives of the enriched microbiome, RS-HXMU1420-13-MAG07, WTSP-JAKVCT01-MAG08 and RS-CAXQZM01-MAG09 (**Figure 3D**). They shared eight MAGs with the NATL, particularly the mid and Eastern regions, where more varied *Trichodesmium* subclade diversity was observed (**Figure 3E**). This group also included enriched microbiome representatives NATL-HXMU1420-13-MAG14, and NATL-HXMU1420-13-MAG02 and NATL-CAXQZM01-MAG17 (**Figure 3E**). MAG NATL-CAXQZM01-MAG17 was detected in all regions and associated with *Trichodesmium* populations characterized by ThiebD, ThiebC subclades, and TeryA-ThiebB codominant samples (**Figure 3E**).

We used indicator species analysis (ISA) and linear correlations between *Trichodesmium* subclade-specific genes and heterotroph MAG percent genome detection to identify associations between heterotrophs and *Trichodesmium* populations (**Figure 3B-E**). The built-in flexibility of the ISA method allowed us to group the detection of specific heterotrophic MAGs by either location or *Trichodesmium* subclade present.

Statistically supported relationships were observed in the NATL. NATL-G030657355-MAG24, NATL-JANSYL01-MAG07, NATL-JANSYL01-MAG01, NATL- JAKVCT01-MAG01, NATL-Rhodobacter_B-MAG01, NATL-UBA2562-MAG01, NATL-Nisaea-MAG01, and NATL-Microscilla-MAG10 were strongly associated with ThiebC- and ThiebD-dominated *Trichodesmium* populations (**Figure 3B**). Five of these MAGs (NATL-G030657355-MAG24, NATL-JANSYL01-MAG07, NATL-JANSYL01-MAG01, NATL-Nisaea-MAG01, and NATL-Microscilla-MAG10) were also associated with the NATL when grouped by location (**Figure S7**). ThiebD abundances positively correlated with the percent genome detection of 12 NATL MAGs (**Figure 4A**) that included all MAGs in the shared NATL microbiome except NATL-G030657355-MAG24 (**Figure 3B**). Moving eastward across the NATL, *Trichodesmium* populations shifted into more mixed communities. This transition also coincided with the loss of several Thieb-associated heterotrophs including NATL-Nisaea-MAG01 and NATL-Microscillaceae-MAG10 (**Figure 3B**).

**Figure 4.**
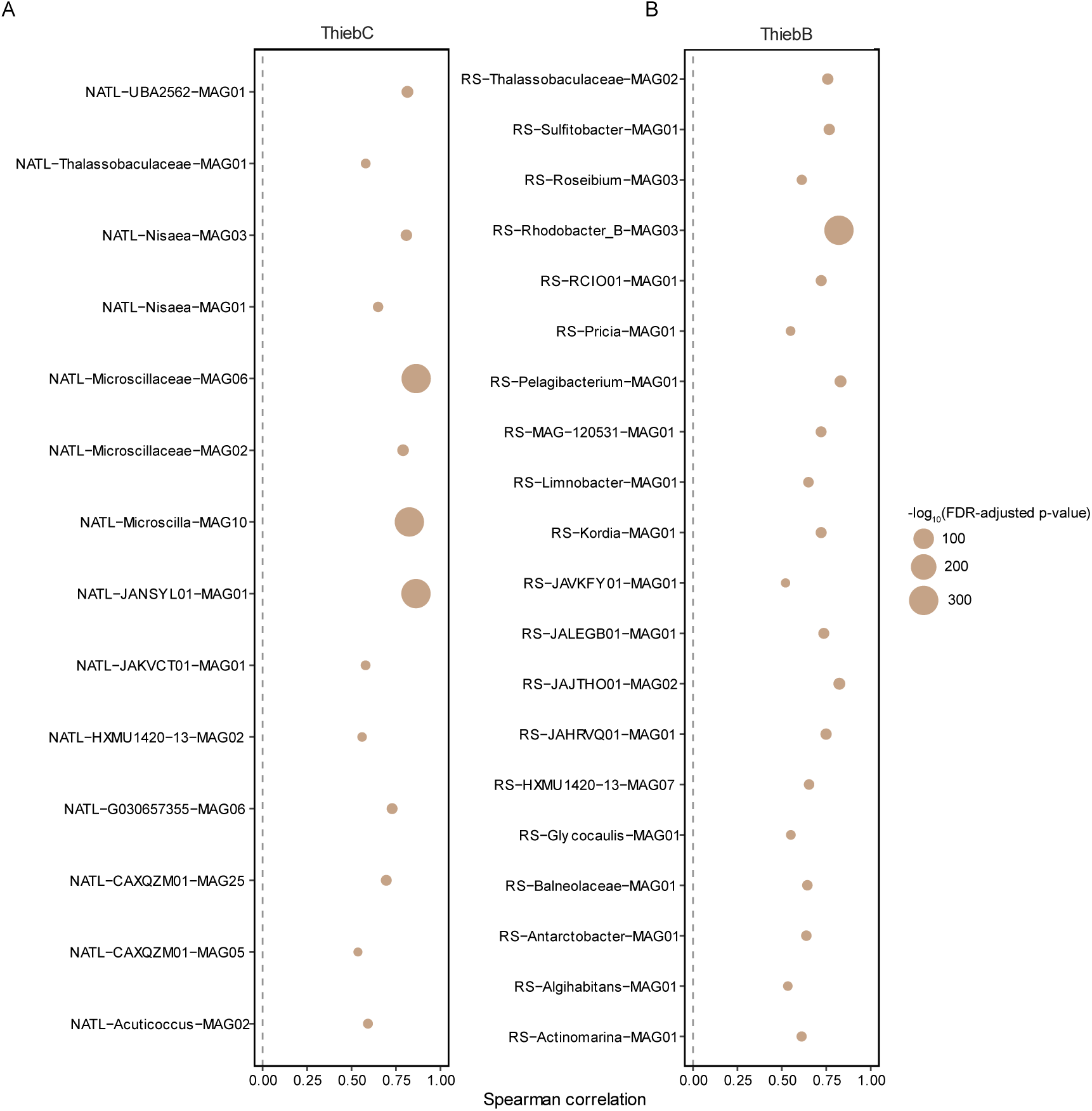
Linear correlations of *Trichodesmium* subclades **(A)** ThiebD or **(B)** ThiebB with heterotrophs. *Trichodesmium* mean coverage (abundance) was correlated with the percent of each heterotroph genome detected. Circles indicate MAGs in the NATL microbiome. Squares indicate MAGs in the putative RS microbiome. Only correlations with spearman rank values > 0.5 are shown. FDR-adjusted p-values were -log10 transformed.

The RS *Trichodesmium* microbiomes differed from those in the NATL, and exhibited several significant *Trichodesmium-*heterotroph associations. Based on the ISA, RS-Sulfitobacter-MAG01 and RS-HXMU1420-13-MAG07 were associated with ThiebB- dominated and ThiebB-TeryA co-dominant populations (**Figure 3C-D**). Both MAGs are also correlated with ThiebB abundance (**Figure 4B**). When grouped by location, 22 MAGs were significantly associated with the RS (**Figure S7**), 11 of which were also correlated with ThiebB (**Figure 4B**). Unlike the NATL, where a shared set of heterotrophs were consistently detected, RS heterotroph and *Trichodesmium* communities varied temporally (**Figure 3D**), with the TeryB-dominated sample supporting a different microbiome relative to the ThiebB-dominant samples (**Figure 3D**).

Importantly, none of the 83 representative MAGs were detected in the St17 bulk plankton tow (**Figure 3B-D**). We also tested for the presence of common phytoplankton-and particle-associated heterotrophs. For example, no *Alteromonas spp.* assembled in the metagenomes collected for this study, and *Alteromonas macleodii* was only detected sporadically through read recruitment (**Table S8**).

### 4.4 Putative Lifestyle Traits and Metabolic Potential of *Trichodesmium*-Enriched Heterotrophs

We investigated motility, genome size, predicted doubling time, and GC% in the enriched microbiome MAGs (**Figure 5**; **Table S9**) because these traits can correlate with growth rates, catabolic niches, and interaction potential with phytoplankton (69,78,79). Motility, inferred from the presence of the MotA/MotB chemotaxis proteins and genes for flagella biosynthesis, was detected in approximately half of the enriched groups (75)(**Figure 5; Table S10**). Genome sizes varied substantially among taxa, ranging from 1.8 to 9.0 Mbp. *Microscilla* had the largest genome (9.0 ± 0.830 Mbp), followed by *Microscillaceae* (4.68 ± 0.51 Mbp), and *Rhodobacter_B* (4.42 ± 0.50 Mbp), while the remaining groups ranged from 1.8–3.2 Mbp (**Figure 5A**). Codon usage bias was used to predict minimum doubling times in hours, with doubling times ≥ 5 hours indicating slow growth (69). The *Microscilla* and *Microscillaceae* groups have predicted doubling times of 4.5 ± 0.4 and 4.9 ± 0.58 hours, respectively, while the rest of the doubling times were predicted to be ≥ 5 hours (**Figure 5B**). GC content separated the MAGs into high GC% (average 59.64%) and low (average 37.73%) content groups (**Figure 5C**)

**Figure 5.**
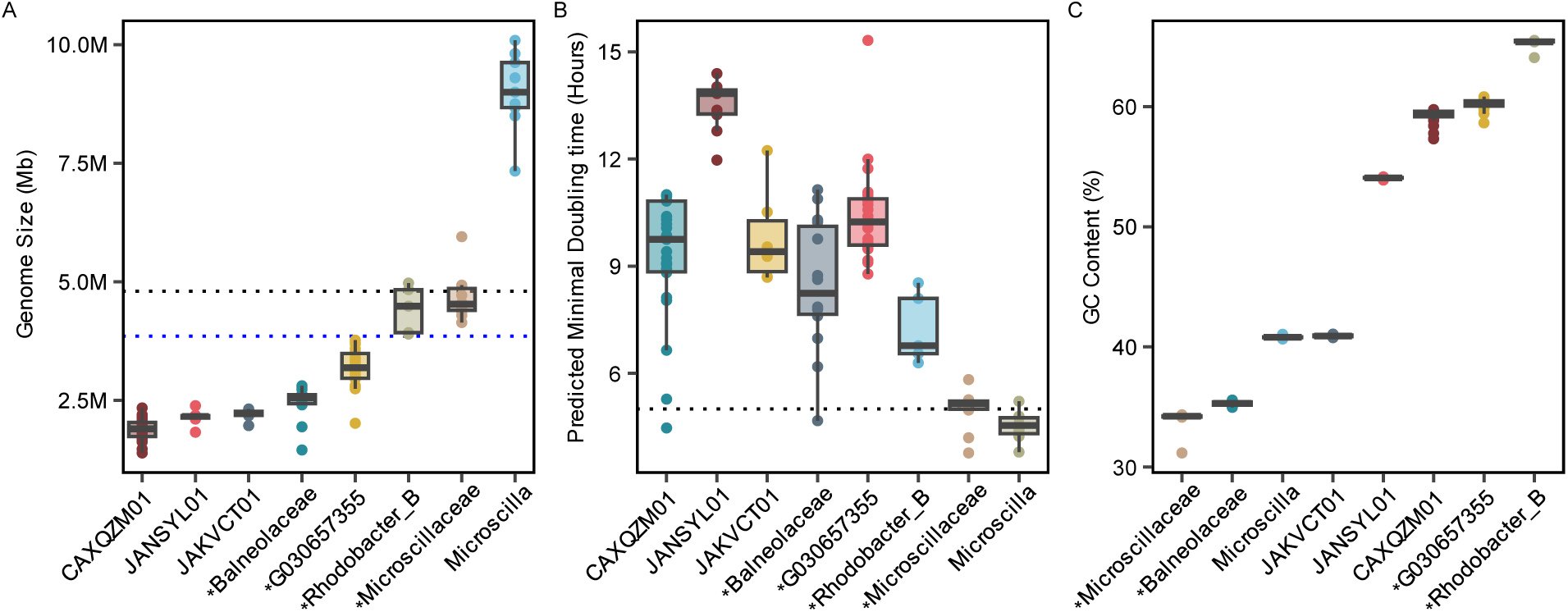
Genome statistics of the *Trichodesmium*-enriched heterotrophs. Groups were included if they had > 5 MAGs in each classification. MAGs were ≥ 70% complete and had < 5% contamination. Genome size and GC content are based on CheckM outputs. **(A)** Genome size. Black line indicates the genome size of a copiotroph, blue line indicates the genome size of an oligotroph defined in Lauro et al 2007. Asterisks denote motile MAGs **(B)** Predicted doubling times based on codon usage statistics and **(C)** GC content.

To connect genome characteristics to potential interactions with *Trichodesmium*, we surveyed 63 complete KEGG functions in 17 representatives of the enriched epibionts. Our analyses focused on catabolism, photoheterotrophy, and phosphorus, nitrogen and iron cycling (**Figure 6**). Multi-enzyme pathways were considered present with ≥ 70% completion scores (**Table S10**).

**Figure 6.**
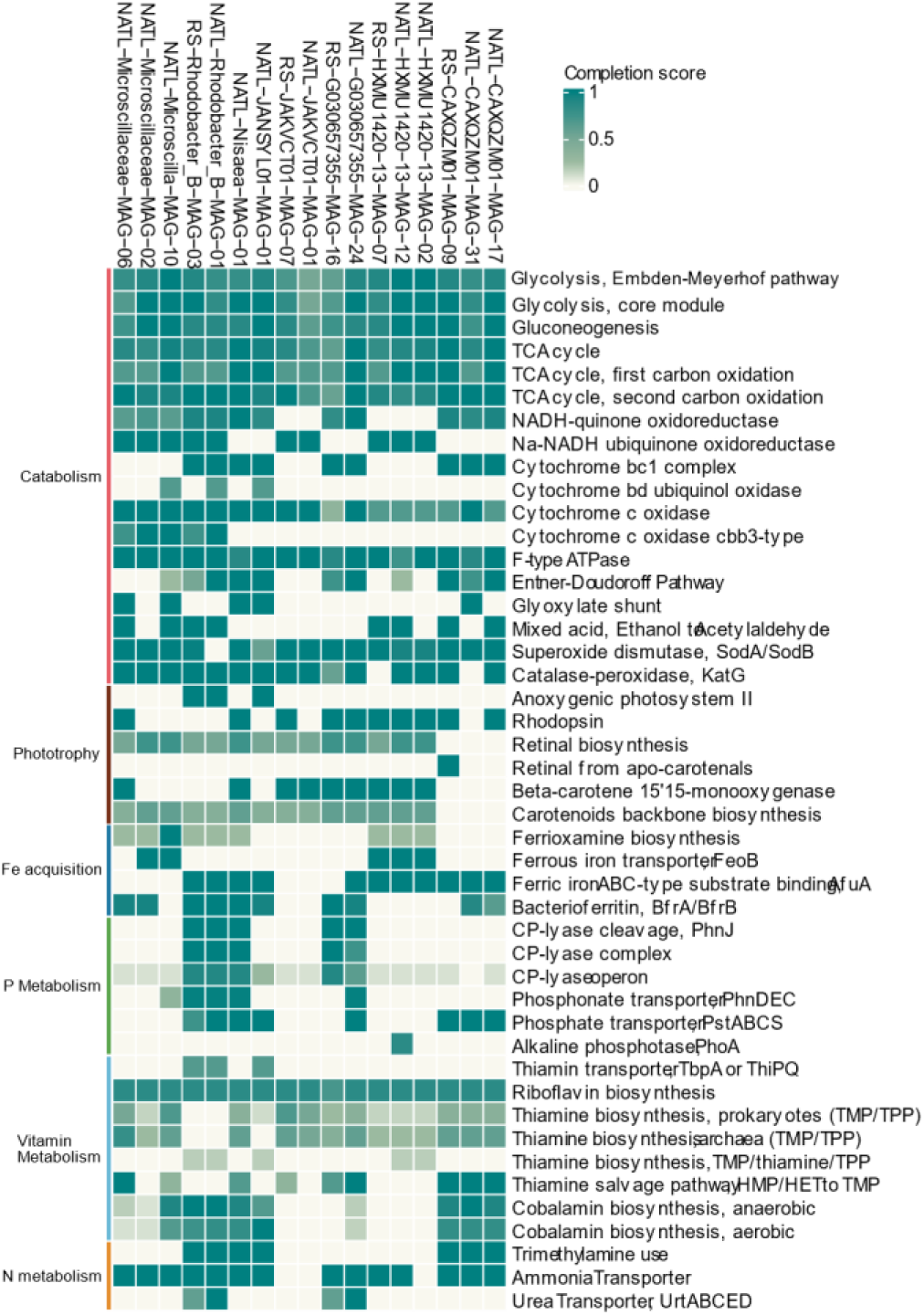
Predicted metabolic pathways of MAGs enriched in the Red Sea and NATL. MAGs were refined and annotated using prodigal and KEGG Orthology. Mutli-enzyme pathways were considered complete if ≥ 70% of the genes were present in a MAG. The best representative MAG from each group was chosen

#### Catabolism and Photoheterotrophy

While all of the MAGs encode for the aerobic cytochrome c oxidase, the *Rhodobacter_B*, and *Microscilla* groups also encode for the low-oxygen cytochrome c oxidase cbb3 (80)(**Figure 6**). To mitigate oxidative stress, the majority of the representative MAGs encode for superoxide dismutase (*SodA* and/or *SodB*) and catalase-peroxidase (*KatG)*. All MAGs are also predicted to perform glucogenesis, synthesizing glucose out of non-carbohydrate sources such as amino acids. Thirteen MAGs (∼81%) are predicted photoheterotrophs using either an anoxygenic type II reaction center or proteorhodopsin. The latter MAGs likely synthesize retinal from apo-carotenals and beta-carotene-15,15’-monooxygenase (81,82) and based on conserved residues at position 105 are predicted to encode “green-light” proteorhodopsin (83).

#### Iron metabolism

Only NATL-Microscilla-MAG10 encoded a complete ferrioxamine siderophore biosynthesis pathway (**Figure 6**). Most MAGs (64%) encoded ferric iron uptake via ABC-type transporters (84), whereas members of the *Microscilla* and HXMU1420-13 groups encoded the ferrous iron transporter *FeoB*. Approximately half of the representative MAGs also encoded the iron storage protein bacterioferritin.

#### Phosphorus metabolism

The enriched MAGs encoded pathways for the acquisition of inorganic and organic phosphorus (**Figure 6**). Four MAGs contained both phosphonate transporters and a complete CP-lyase pathway, enabling the use of alternative sources of organic P during depletion (14). Most MAGs also encode transporters for phosphate and/or phosphonate uptake. Among the representative genomes, only NATL-HXMU120- 13-MAG-12 encoded the alkaline phosphatase *PhoA,* which breaks down C-O-P bound organic phosphorus (85).

#### Vitamin Metabolism

Consistent with previous studies showing that *Trichodesmium* is a de novo producer of B-vitamins in the oceans (86,87), many of the enriched MAGs are predicted to be thiamin (B_1_) and cobalamin (B_12_) auxotrophs.

#### Nitrogen metabolism

Three complete nitrogen use pathways were predicted in the enriched MAGs (**Figure 6**). Although most MAGs encode ammonia transporters, four also encode urea transporters. The potential to use trimethylamine use as an alternative organic C & N source (40) was predicted in seven MAGs based on the presence of the *mttB2* gene and dimethylamine/trimethylamine dehydrogenase genes.

## 5. Discussion

We investigated the genomic characteristics of the *Trichodesmium*-associated microbiome and the processes governing heterotroph community assembly across major ocean basins. Although previous studies established the existence of *Trichodesmium’s* microbiome (25,26), the ecological processes shaping its composition and the relationship between microbiome structure and *Trichodesmium* subclade diversity have remained largely unexplored.

By combining newly generated BATS metagenomes with published datasets from the NATL, WTSP, and RS, we identified heterotrophic MAGs repeatedly recovered across samples, years, and ocean basins, indicating persistent associations with *Trichodesmium* colonies. Across regions, microbiome composition was structured by geography and *Trichodesmium* composition, with deterministic processes dominating microbiome assembly. Consistent with this pattern, stochasticity analyses indicated that deterministic processes play a dominant role in shaping microbiome assembly. Together, these findings suggest that *Trichodesmium* colonies harbor predictable microbial assemblages shaped by both host population structure and environmental context.

### 5.1 *Trichodesmium* colonies are associated with a unique heterotrophic community

*Trichodesmium* colonies harbor a phylogenetically and ecologically unique microbiome. The large number of novel MAGs identified by GTDB-tk (77), indicates that many assembled taxa are underrepresented in existing oceanic and phytoplankton-associated datasets and may be rare in the oligotrophic ocean (**Table S4**). One example is the CAXQZM01 lineage, a phylogenetically distinct and biologically novel group (**Figure 1**). Members of this lineage have been isolated from *Trichodesmium* cultures (GenBank JN790965.1; JN790966.1) and consistently recovered from field samples across regions and years, indicating a persistent association (**Figure 1**).

In contrast, *Alteromonas*, a common phytoplankton-associated heterotroph (88–91), was only detected sporadically and was not a consistent member of the microbiome. Although *Alteromonas* has previously been detected on colonies (25,41,92), its occurrence may be transient, potentially reflecting colonization during senescence or other stressful events (93). Moreover, none of the core microbiome members were detected in the bulk plankton tow sample (**Figure 3**), suggesting they are not broadly associated with particles or other phytoplankton. Together, these observations support the existence of a *Trichodesmium*-specific microbiome composed of both novel and persistent bacterial lineages.

Genomic characteristics of the enriched microbiome suggest multiple ecological strategies coexist within *Trichodesmium* colonies (**Figure 5**). Approximately half of the representative MAGs had small genomes, limited mobility potential, and slow predicted growth rates, traits characteristic of oligotrophic heterotrophs (69,78). The remaining MAGs exhibited features more typical of ecological opportunists, including larger genomes and greater metabolic versatility (78,94–96). Notably, roughly half also had elevated GC content (**Figure 5**), a trait that has been linked to greater N availability (97), suggesting sustained accesses to N despite the surrounding nutrient-limited waters.

One explanation is that *Trichodesmium*’s slow growth rate (98) limits resource turnover, favoring recruitment of rarer oligotrophs, while also allowing copiotrophic taxa to persist. Similar patterns have been observed in freshwater cyanobacteria (99). These findings contrast with the view that phytoplankton-microbiomes are dominated by fast-growing, motile copiotrophs (95,100,101), instead suggest that *Trichodesmium* colonies support microbial communities spanning a range of ecological strategies.

The enriched heterotrophic community in the NATL was dominated by a small number of taxa repeatedly recovered across samples and locations (**Figure 1**). Their consistent recovery suggests they are physically attached to *Trichodesmium* filaments and vertically transmitted. These patterns indicate that colonies consistently recruit or maintain the same community. Although some enriched taxa were shared among ocean basins, overlap was limited and community composition remained regionally distinct (**Figures 2 & 3**).

Several enriched taxa comprised multiple closely related but genetically distinct lineages. For example, *Microscilla* MAGs clustered into three phylogenetic groups, including two novel lineages. *Microscilla* and related *Microscillaceae* MAGs were associated with ThiebD populations and declined as ThiebC became dominant. Consistent with previous observations in the NATL (26), these results identify *Microscilla* as a recurrent member of the *Trichodesmium* microbiome represented by multiple, closely related species. Similarly, HXMU1420-13 MAGs previously detected in RS colonies (25), formed distinct lineages with different distributions across regions and *Trichodesmium* populations. Together, phylogenomic placement and read recruitment patterns indicate that enriched heterotrophs are structured by *Trichodesmium* subclades, which are themselves associated with ocean basin (**Figures 1, 2 & 3**).

### 5.2 Different *Trichodesmium* Populations Support a Unique Microbiome Across Space and Time

*Trichodesmium* populations were consistent over space and time, with distinct subclades enriched in different ocean basins (**Figures 2 & 3**). Although *Trichodesmium* abundance is variable, population structure is highly correlated with regional biogeochemical regimes (34,35,102). ThiebD dominated the western NATL over a decade, suggesting a resident population (**Figures 2 & 3**). In contrast, mid-Atlantic colonies contained mixed ThiebD-ThiebC populations, whereas ThiebC dominated the eastern NATL (**Figures 3**). These patterns parallel nutrient gradients across the NATL, with P limitation in the west, co-limitation in the mid-Atlantic, and Fe limitation in the east, all which influence *Trichodesmium* abundances and activity (9,17,102). Similar regional consistency was observed elsewhere, with ThiebB dominating the RS across multiple sampling periods and TeryA-ThiebB co-dominating the WTSP (**Figure 3**), although the latter was supported by fewer samples. While disentangling subclade identity from environmental influences remains difficult, these results demonstrate persistent regional enrichment of specific *Trichodesmium* populations across ocean basins.

NMDS clustering captured the specificity and stability of the heterotrophic community (**Figure 2**). Within the NATL, western and eastern communities formed distinct clusters, while mid-Atlantic samples occupied an intermediate position, reflecting their mixed *Trichodesmium* composition. Across the global dataset, NATL samples remained clustered together despite spanning a decade of sampling, indicating long-term stability. Similarly, BATS colonies showed no significant interannual or seasonal shifts in microbiome composition, despite the region’s strong seasonality (103). These observations suggest that nutrient fluctuations have limited influence on core the heterotrophic community, and that persistent ecological associations promote community stability.

Patterns of community overlap further support a link between microbiome composition and *Trichodesmium* population structure. For example, NATL and WTSP colonies shared several heterotrophic taxa despite their geographic separation, potentially reflecting the occurrence of shared Thieb subclades (**Figure 2 & 3**), whereas overlap between the RS and WTSP may be driven by TeryA populations (**Figure 2 & 3**). Because these heterotrophs remained tightly associated with colonies, positive correlations with specific *Trichodesmium* subclades (**Figure 4**) likely reflect direct interactions rather than oceanographic conditions alone. Overall, this consistency combined with the low normalized stochasticity ratios indicates low community turnover, which is congruent with deterministic factors shaping the microbiome (104,105).

Read-recruitment and indicator species analyses identified putative ThiebD and ThiebC core microbiomes (**Figures 3 & 4**). In the NATL, NATL-JAKVCT01-MAG01, NATL-Microscilla_MAG10, and NATL-Nisaea_MAG01 were strongly associated with these subclades and consistently co-occurred with Thieb colonies across sampling periods. While other subclades were represented by fewer samples, additional patterns support subclade-specific microbiome structure. For example, ThiebD- and ThiebC- enriched colonies collected at the same time harbored distinct heterotroph communities, and in the RS, TeryB communities differed from co-occurring ThiebB communities. Given the strong dispersal limitation and regional selection characteristic of marine microbial communities (104–106), shared heterotrophs across geographically separated regions may reflect similar selective pressures associated with *Trichodesmium* subclade traits, such as physiology or morphology. Although the dominance of ThiebD limits our ability to fully disentangle subclade and environmental effects, shifts in heterotroph occurrence corresponding to changes in *Trichodesmium* populations (**Figure 3**) support subclade-specific microbiome recruitment.

MAGs detected in multiple *Trichodesmium* populations may represent microbiome generalists that associate with multiple subclades (**Figure 3E**). In contrast, intermittently detected members likely comprise an episodic community (**Figure 3F**), with occurrence influenced by *Trichodesmium* physiology and environmental conditions. Based on this, *Trichodesmium* hosts a stable core microbiome alongside a dynamic community shaped by local environmental conditions.

Parallel patterns in microbiome composition and *Trichodesmium* population structure are consistent with deterministic structuring, although the underlying drivers cannot be fully resolved. While *Trichodesmium-*mediated filtering is likely important, contributions from abiotic factors and stochastic forces cannot be excluded. Shared environmental preferences, differences in sampling methods among studies, and an incomplete understanding of *Trichodesmium* biogeography complicate comparisons across regions. In other phytoplankton microbiomes, stochastic processes can strongly influence microbiome assembly or modulate the effects of selection (28,30). However, colonies with *Trichodesmium* populations but collected from the same water mass harbored distinct microbiomes, a pattern difficult to reconcile with neutral processes alone. Consistent with this, normalized stochasticity ratios indicate a strong role for deterministic processes in the NATL microbiome (**Table S6**). Although additional work is needed beyond the NATL, particularly in regions where *Trichodesmium* is an important N fixer, these results suggest that *Trichodesmium* subclade diversity is a particularly strong driver of microbiome composition.

### 5.3 Characteristics of *Trichodesmium* Microbiome

Predicted carbon metabolism, low-oxygen cytochrome c oxidases, and other anaerobic pathways (**Figure 6**), indicate that these heterotrophic MAGs are facultative aerobes capable of fermentation within the rapidly fluctuating O_2_, CO_2_, and pH gradients in colonies (107–109). The consistent detection of green-light proteorhodopsins and anoxygenic photoheterotrophy further suggests that some epibionts use light harvesting systems adapted to organic matter-rich water (110,111), reducing competition with *Trichodesmium* for blue wavelengths characteristic of the oligotrophic ocean. The complex colony geometries likely creates diverse micro-environments that support metabolic diversity, reduce niche overlap, and promote the coexistence of novel and closely related lineages (**Figure 1 & Table S4**).

Predicted metabolic capabilities suggest mutually beneficial host-microbiome interactions in P, N, Fe and vitamin-related processes within *Trichodesmium* colonies (**Figure 6**). For example, heterotrophs encoding CP-lyase pathways may enhance access to organic P, thereby alleviating P-limitation in colonies. Likewise, heterotrophic trimethylamine use may reduce the energetic cost of N fixation, as trimethylamine inhibits N fixation without affecting growth (112). Conversely, while epibionts have been implicated in Fe acquisition (21,25), siderophore biosynthesis was predicted in only one MAG (**Figure 6**), highlighting the need to quantify the microbiome contributions to in Fe acquisition across oceanographic contexts. Similarly, while bacteria are key facilitators of B_1_ and B_12_ production in phytoplankton-microbiomes (29,113), predicted auxotrophies among enriched *Trichodesmium* epibionts suggest the opposite relationship, with *Trichodesmium* producing B_12_ and other B-vitamins for the microbiome (86,87). Future ‘omics studies will be needed to determine how community members partition these functions and contribute to nutrient cycling across *Trichodesmium* populations.

### 5.4 Conclusions and Future Directions

We characterized the *Trichodesmium*-microbiome in the context of subclade identity, showing that subclades may structure epibiont communities. Our findings suggest that distinct biogeochemical regimes across ocean basins select for specific *Trichodesmium* populations, which in turn shape microbiome composition and stability. These results indicate that the microbiome may contribute to *Trichodesmium* biogeography, abundances, and physiology. By establishing the stability and specificity of the *Trichodesmium* microbiome, this work provides a framework for future studies of host-microbiome interactions. A more complete understanding of the processes governing colony diversity, abundances, and microbiome composition will improve predictions of N_2_ fixation and primary productivity in a changing ocean.

## Acknowledgements

We thank the scientific staff and crew of the R/V Atlantic Explorer and of the AT39-05/Tricolim research expedition.

